# Lactate production without hypoxia in skeletal muscle during electrical cycling: Crossover study of femoral venous-arterial differences in healthy volunteers

**DOI:** 10.1101/355321

**Authors:** Jan Gojda, Petr Waldauf, Natália Hrušková, Barbora Blahutová, Adéla Krajčová, Petr Tůma, Kamila Řasová, František Duška

**Author notes:** JG and PW contributed equally and shall be considered joint first authors. FD is a senior author. Corresponding author (JG).

## Abstract

**Background:** Aim of the study was to compare metabolic response of leg skeletal muscle during functional electrical stimulation-driven unloaded cycling (FES) to that seen during volitional supine cycling.

**Methods:** Fourteen healthy volunteers were exposed in random order to supine cycling, either volitional (10-25-50 W, 10 min) or FES assisted (unloaded, 10 min) in a crossover design. Whole body and leg muscle metabolism were assessed by indirect calorimetry with concomitant repeated measurements of femoral venous-arterial differences of blood gases, glucose, lactate and amino acids.

**Results:** Unloaded FES cycling, but not volitional exercise, led to a significant increase in across-leg lactate production (from −1.1±2.1 to 5.5±7.4 mmol/min, p<0.001) and mild elevation of arterial lactate (from 1.8±0.7 to 2.5±0.8 mM). This occurred without widening of across-leg VA O_2_ and CO_2_ gaps. Femoral SvO_2_ difference was directly proportional to VA difference of lactate (R^2^= 0.60, p=0.002). Across-leg glucose uptake did not change with either type of exercise. Systemic oxygen consumption increased with FES cycling to similarly to 25W volitional exercise (138±29% resp. 124±23% of baseline). There was a net uptake of branched-chain amino acids and net release of Alanine from skeletal muscle, which were unaltered by either type of exercise.

**Conclusions:** Unloaded FES cycling, but not volitional exercise causes significant lactate production without hypoxia in skeletal muscle. This phenomenon can be significant in vulnerable patients’ groups.

## Introduction

Functional electrical stimulation-assisted cycling (FES cycling) is a method originally developed over 30 years ago for patients with spinal cord injury [1]. It uses computer-driven electrical pulses delivered by transcutaneous electrodes and directly activating muscle contractions, independently on functionality of the physiological pathway between upper motoneuron and the neuromuscular junctions. The method is now commercially available in the form of both stationary and mobile devices [2], used by patients with a wide range of conditions incl. spinal cord injury [3], stroke [4,5], and multiple sclerosis [6]. FES cycling was demonstrated to improve cardiovascular fitness, insulin sensitivity [7] bone density and muscle strength [2,8]. In recent years, FES-cycling has become particularly attractive for sedated critically ill patients. Early mobilization is the only intervention, which can partially prevent the development of intensive care unit-acquired weakness [9–14] - the major long-term consequence in the survivors of protracted critical illness [15,16]. Muscle atrophy [17,18] and dysfunction [18] occur very early in the critically ill and FES cycling can help to deliver exercise before the patient can co-operate with a physiotherapist [19].

Although FES cycling seems to be feasible in intensive care unit patients [19], before its effect on meaningful clinical outcomes can be tested in the critically ill and other vulnerable patients groups, important physiological questions need to be addressed. Metabolic efficacy (i.e. power output divided by metabolic cost) of the FES cycling is typically very low, around 5-10%, as compared to 25-40% in volitional cycling [20–22]).

This is likely due to non-physiological pattern of muscle activation, where large muscle groups are activated simultaneously rather than small well-coordinated units [2,23]. Despite FES cycling increases cardiac output [24] and leg blood flow to the same extent [25] or even more [26] than volitional cycling and consequently oxygen delivery to the muscle should be normal, there are features suggesting early switch to anaerobic metabolism: early fatigue [23,27], rapid intramyocellular glycogen depletion [28], increase of respiratory quotient (RQ) >1 [20] and even a mild increase in arterial lactate levels [29]. Nonetheless, a direct evidence of the presence of anaerobic metabolism in skeletal muscle during FES cycling is lacking. Moreover, whilst the influence of volitional resistance exercise on amino acid metabolism has been extensively studied [30–34] there is no such data for FES cycling. These questions may be particularly relevant before FES-assisted exercise is introduced to critically ill patients, who are in profound protein catabolism and may be less able to clear lactate from systemic circulation.

In light of this we conducted a crossover study of volitional and FES supine cycling in healthy postprandial volunteers, where we combined indirect calorimetry with across-leg VA difference studies. We hypothesized that FES-cycling as compared to light volitional exercise would lead to increased production of lactate in correlation with widening of VA-CO_2_ gap (as the measure of anaerobic metabolism), and with increased amino-acid efflux from skeletal muscle during exercise.

## Materials and methods

### Overview of study design

The study was performed during two visits performed 1 week apart. Subjects were asked to attend the visit at 08:00 AM after an overnight fast. In between these visits, the subjects were advised to take their usual diet and avoid strenuous exercise. During the first visit, the volunteers underwent a physical examination and body composition measurement. After 30 min bed rest, their energy expenditure was measured using indirect calorimetry with a ventilated canopy system. Afterwards, in each subject’s VO_2MAX_ was determined on a cycle ergometer with stepwise load by 25 W increments until exhaustion. During the second visit, subjects were given a standardized breakfast containing 70 g of carbohydrates, 10 g protein and 15 g of fat. Afterwards, femoral vein and radial artery were cannulated. After 30 min rest, the subjects were exposed in random order to one of two supine exercise protocols, separated by 3 hours rest. Both protocols begun with baseline measurements (AV difference studies and calorimetry) followed by 5 min of passive cycling. Then, the subjects either performed three 10 min cycles of volitional cycling (at 10, 25 and 50 W, respectively) separated by 5 min of passive cycling (Group A), or FES cycling (Group B). The exercise protocols are outlined in Figure 1. University Hospital Kralovske Vinohrady’s Ethical Review Board reviewed the protocol and approved the study. Prior to the enrolment, all subjects gave their written informed consent in accordance with the Declaration of Helsinki.

**Fig 1.**
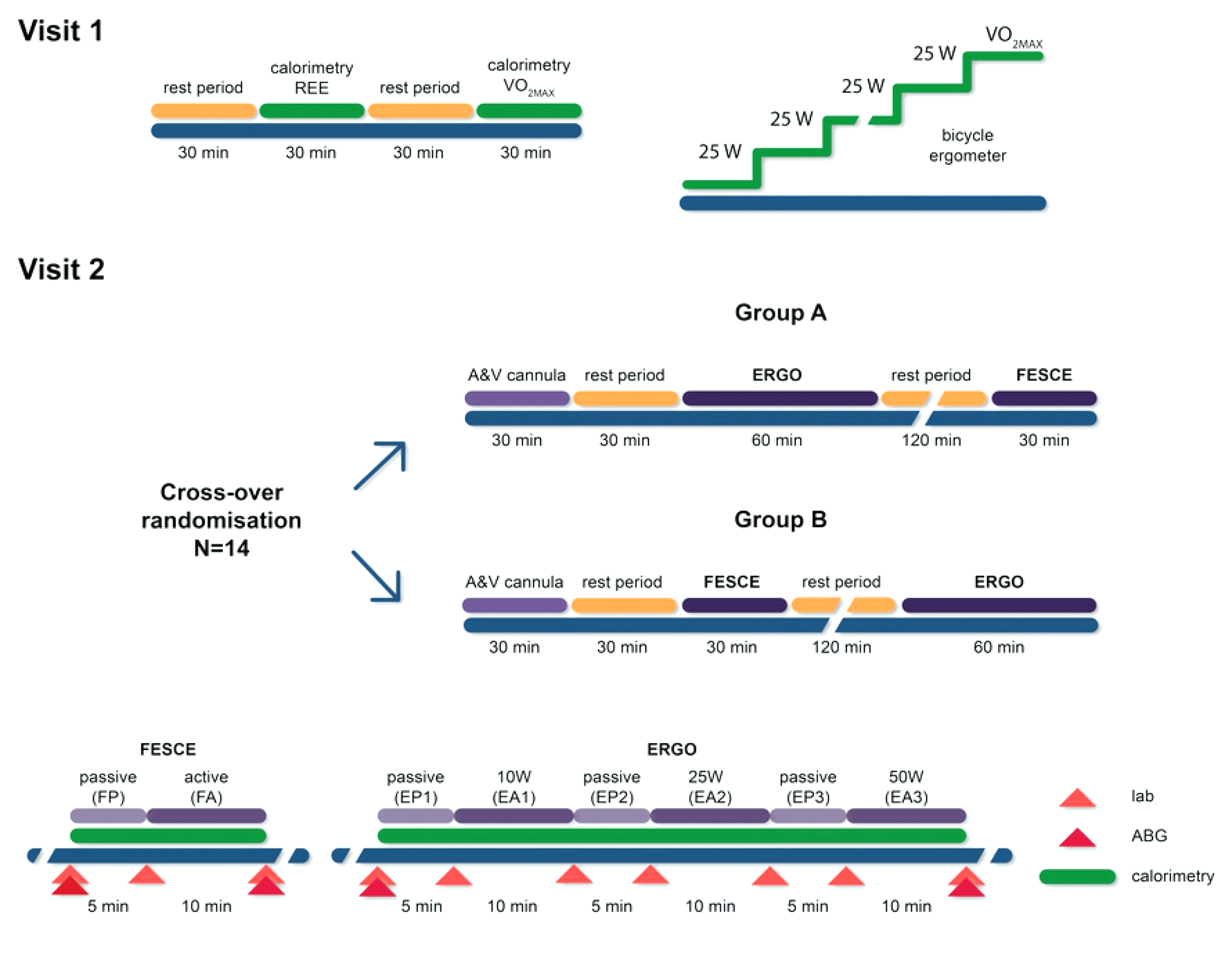
Overview of study design. Arrows designate arterial and venous blood sampling times. Note: ERGO = volitional cycling, FESCE = functional electrical stimulation cycling. Details of exercise are shown in the inlet at the bottom.

### Study subjects

Our experimental group consisted of 14 healthy volunteers. Their baseline characteristics are given in Table 1. Body fat was assessed using bioimpedance analysis (NutriGuard 2000, Bodystat, Germany).

**Table 1.** Baseline characteristics of study subjects.

| Parameter | Mean±SD | N |
| --- | --- | --- |
| Age (years) | 31±8 | 14 |
| Sex (M/F) | 11/3 | 14 |
| BMI (kg/m<sup>2</sup>) | 23.7±3.7 | 14 |
| Body fat (%) | 14±6 | 14 |
| REE (kcal/day) | 1901±356 | 13 |
| RQ at rest | 0.90±0.10 | 13 |
| VO<sub>2MAX</sub> (ml/kg.min) | 41±6 | 13 |

Note: BMI = body mass index, REE = resting energy expenditure, RQ = respiratory quotient.

## Methods

### Indirect calorimetry

Resting energy expenditure and RQ were measured after overnight (12 h) fast and 30 min bedrest using canopy Quark RMR device (Cosmed, Italy). To determine peak oxygen uptake (VO_2max_) exhaustive exercise test was performed in each subject on an electromagnetically braked bicycle ergometer Ergoline Ebike (Ergoline Gmbh, Germany). After 5 min warm-up period, a workload of 50W was initiated and increased by 25 W every minute continuously until fatigue despite the verbal encouragement. Oxygen uptake was measured using Quark RMR device (Cosmed, Italy). ECG was monitored continuously.

### Cannulations

Femoral vein was cannulated 2-3 cm below inguinal ligament under ultrasound guidance. In order to avoid the admixture of blood from saphenous and pelvic veins [35], a single-lumen central venous catheter (B-Braun, Germany) was inserted retrogradely to the depth of 10-15 cm so that the tip was deep in the femoral muscular compartment. For arterial sampling, we used a 22 F catheter (BBraun, Germany) inserted into the radial artery.

For both volitional and FES cycling we used RT-300 bikes (Restorative Therapies Ltd., USA) and the exercise was performed in supine position. *Volitional cycling* consisted of three 10 min intervals of active cycling: 10W (13 revolutions/min, resistance 7 N/m), 25W (31 revolutions/minute, 7.6 N/m), 50W (35 revolutions/min, and resistance 13.4 N/m). These period were preceded (warm up) and separated by 5 min of passive cycling at revolutions/min. *FES cycling*: Three pairs of transcutaneous electrodes (3 × 4″, Restorative Therapies, Ltd., USA) electrodes were applied on each leg over quadriceps, hamstrings and gluteus maximus muscles, as per manufacturer’s instructions. Prior to electrode placement, we measured the thickness of fat layer between the skin and muscle by ultrasound. After 5 min passive warm up (25 revolutions/min), the target speed was changed to 30 revolutions/min and stimulation gradually (1%/s) started to achieve 25 mA. Then, in each subject, the stimulation current was gradually increased to reach subjectively tolerated maximum. Oxygen uptake was measured continuously in both volitional and FES assisted cycling using mask breath-by-breath system (Quark RMR device, Cosmed, Italy).

### Laboratory methods

Arterial and venous blood samples were analysed for blood gases, lactate and haemoglobin using POCT analyser Cobas b221 (Roche Diagnostics Limited, USA). For other analysis blood samples were centrifuged and frozen at −80°C until analysed. Serum creatine kinase and myoglobin was measured in a certified institutional laboratory (Cobas system, Roche Diagnostics Ltd., USA). Serum amino acid concentration in arterial/venous blood was analysed using capillary electrophoresis as described [36].

## Calculations and statistics

### Metabolic efficacy

Metabolic efficacy of volitional cycling was calculated as power output divided by the increase of energy expenditure [2]. Veno-arterial gap in the total content of carbon dioxide (Ct-CO_2_ gap) was calculated according to equations used in ABL 900 Analyser (by Radiometer, Copenhagen, Denmark).
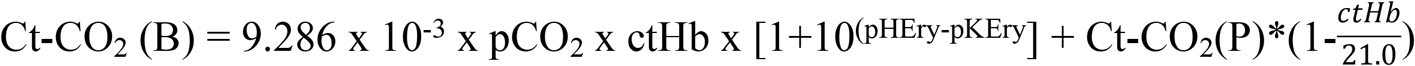

where Ct-CO_2_(B) = CO_2_ content in blood in mmol/L; Ct-CO_2_(P) = CO_2_ content in plasma in mmol/L and equals to 0.23 × pCO2 + cHCO_3_−(P); pCO_2_ is partial pressure in kPa, ctHb = haemoglobin content in mmol/L. Ct-CO_2_(P). pH_ERY_ = estimated intracellular pH in red blood cells, which equals to 7.19+0.77 × (pH-7.4)+0.035 × (1-SO_2_), where S O_2_ is haemoglobin saturation with oxygen; and finally pK_ERY_ is a negative decadic logarithm of bicarbonate dissociation constant = 6.125 −log(1+10^(pHeERY-7-84-0.6*SO2)^).

### Blood flow

In both FES and low intensity volitional cycling, leg oxygen uptake represents a relatively fixed proportion (76±8% vs. 78±9%, respectively) of whole-body oxygen uptake [37]. Therefore, an index of blood flow through the leg was calculated as whole-body oxygen consumption divided by the difference of oxygen content in arterial and femoral-venous blood. Blood oxygen content was calculated in mmol/L as 0.00983*pO_2_ + SO_2_[%]/100 * Hb *0.06206*(1-COHb[%]/100 - metHb[%], where S O_2_ is saturation of haemoglobin with oxygen [%], Hb is haemoglobin [mmol/L], CO-Hb and met-Hb are fractions of carbonyl and methemoglobin, respectively, and pO_2_ is partial pressure of oxygen [kPa]. At rest before volitional and FES cycling, blood flow index was 6.6±2.4 vs. 6.3±3.4 (p=0.57), and increased significantly (p<0.01) and similarly (p=0.77) to 160% and 165% of baseline after volitional and FES exercise.

### Statistics

We used linear mixed effect model for 2×2 crossover design processed with software Stata 15 (Stata Corp., LLC, U.S.A.). The model consists of fixed and random part. In the fixed part, the model contained following parameters: (1) Sequence, i.e. order in which subject performed volitional and FES cycling protocols. Had this parameter been significant, a carry-over effect would have been present; (2) Period, basal vs. active, a parameter exploring the effect of the exercise, regardless whether volitional or FES; (3) Treatment, exploits the difference between volitional and FES cycling; and (4) Interaction Period#Treatment exploits whether FES cycling differs from volitional cycling during exercise period. Random part of the model contains subject number in order to take into account repeated measurements. P value <0.05 was considered as significant. Given the physiological nature of the study, we have not performed a formal power calculation and sample size is based on an assumption that a minimum of 7 subjects would be needed in each group to compare FES-driven and volitional cycling had there been a carry-over effect (i.e. a difference at 2^nd^ baseline caused by previous intervention). In none of the measured values the Sequence parameter was significant (p = 0.14-0.94), so we assume no carry over effect and do not report this value in the Results.

## Results

### Tolerability and signs of muscle damage

All 14 subjects finished the protocol without adverse events; baseline (visit 1) calorimetry data are available for 13 subjects only due to a technical problem. Maximum tolerated stimulation current of FES was 45±13 mA (range 25-67 mA). Although FES cycling caused a degree of discomfort, post-exercise serum myoglobin remained within reference range (<85 ng/mL) in all subjects (33±15 pg/mL, range 21-74). Nonetheless, there was a positive correlation between maximal stimulation current and post-exercise serum myoglobin (R^2^=0.57, p=0.002).

### Metabolic efficacy of volitional vs. FES cycling

Metabolic efficacy of volitional cycling was 39.2±5.6%. Unloaded FES cycling led to an increase of metabolic rate to 138±29% from baseline, which was comparable to the increase with 25 W volitional exercise (124±23%). See Figure 2. Energy gain from anaerobic glycolysis was negligible or negative for volitional cycling and 5.0±6.2 W for FES cycling.

**Fig 2.**
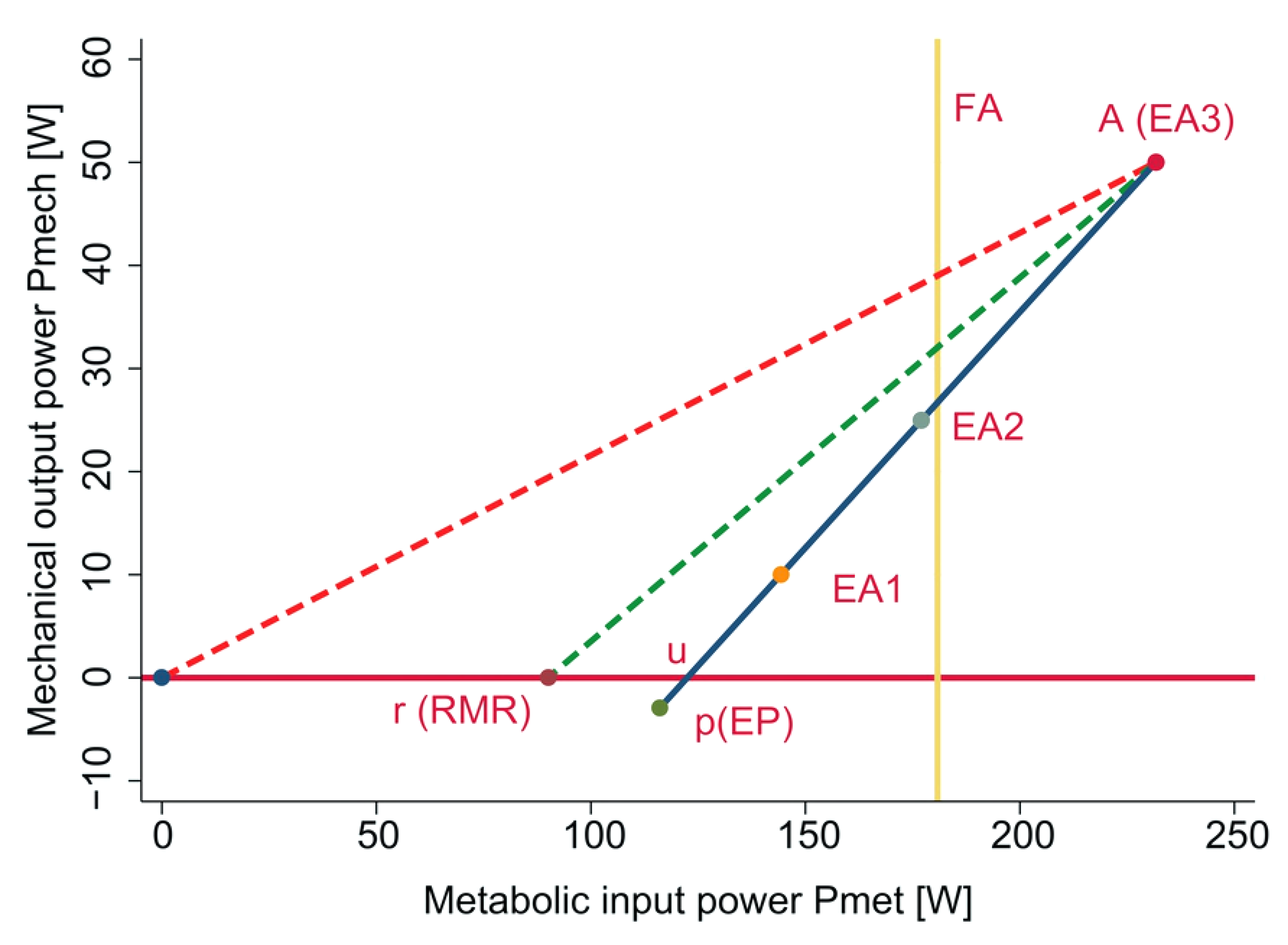
Hunt’s diagram [2] outlining the efficacy of volitional exercise relative to metabolic cost of unloaded FES cycling (yellow line). Note: Metabolic efficiency is the gradient of the line joining the active cycling operating point a to one of the baseline conditions: u is unloaded cycling; r is rest, p is passive cycling.

### Exploring muscle metabolism during FES cycling

With volitional exercise, VA differences of both O_2_ and CO_2_ contents (Ct-O_2_ and Ct-CO_2_) tended to widen with volitional exercise (Fig. 3A and 3B), whilst the opposite trend was seen for FES cycling. In line, there was no change in oxygen saturation of haemoglobin in femoral venous blood neither with volitional exercise (from 63.9±12.7% to 64.3±8.7%), whilst there was a surprising increase after FES cycling (from 62.6±11.3 to 70.3±8.7%; p=0.02). Across-leg respiratory exchange ratio (i.e. the ratio between VA differences of CO_2_ and O_2_ contents) although different at baseline (Fig 3C) tended to increase with volitional cycling, but this change was not significant. There was no change from baseline in across-leg glucose uptake of glucose (FES −5.5±3.9 to −5.9±3.6mmol/min; volitional −7.0±3.6 to −6.9±6.1mmol/min). Whole body RQ increased with FES cycling (0.88±0.02 to 0.95±0.02, p=0.001, but did not change with volitional exercise (0.87±0.02 to 0.85±0.02, p=0.55; See Fig 3D) and only FES cycling led to an increase in across-leg lactate VA differences and production (from −1.1±2.1 to 5.5±7.4 mmol/min, p<0.001 vs. from −0.9±1.1 to −0.4±1.2 mmol/min, p=0.70 Fig 3E) with very high inter-individual variability (See Fig. 3F). Systemic arterial lactate levels remained normal after volitional cycling (from 1.6±0.6 mmol/l to 0.9±2.1 mmol/l, p=0.887), and increased after FES cycling (from 1.6±0.7 mmol/l to 2.3±0.8 mmol/l, p<0.001).

**Fig 3.**
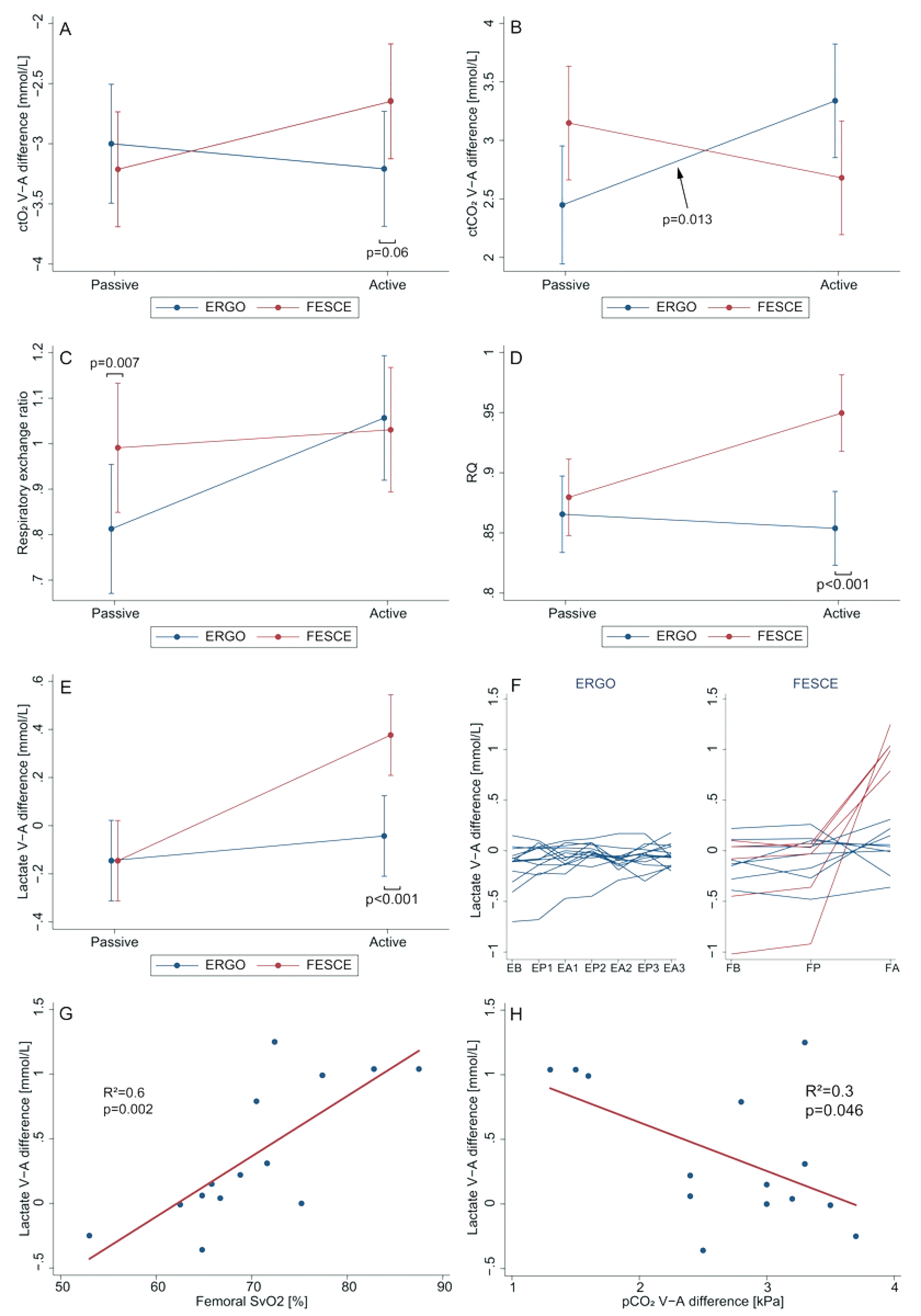
Venous-arterial (VA) differences studies. Lactate VA difference is derived from multiplying femoral VA differences of concentrations and calculated leg blood flow. See text for further details. Note: ctO_2_ and ctCO_2_ = total blood content of oxygen and carbon dioxide; RQ = whole body respiratory quotient; SvO_2_ = femoral venous saturation of haemoglobin with oxygen. ERGO = volitional cycling; FESCE = functional electrical stimulation-assisted cycling; Passive period vs Active FES/50W volitional period.

### Analysing lactate production

There was a significant positive correlation between VA lactate difference and femoral venous haemoglobin saturation with oxygen (R^2^=0.6, p=0.002, Fig 3G) and lactate producers had smaller veno-arterial difference in CO_2_ content of the blood (R^2^=0.3, p=0.046, Fig 3H), effectively ruling out oxygen delivery problem due to low flow. Subjects with femoral VA lactate difference >0.5 mmol/L (“lactate producers”, n=5, see Fig. 3F) were compared with the rest of the group (n=9). Lactate producers tended to have smaller proportion of body fat (8 vs., 13%, p=0.131) and had higher RQ at baseline (0.94±0.06 vs., 0.86±0.07, p=0.034). Of note, stimulation current used during FES cycling was not different in lactate producers (42±10 vs. 44±16 mA, p=0.87).

### Amino acid metabolism

As expected in postprandial volunteers, at baseline resting skeletal muscle was taking up branched-chain amino acids (BCAAs) whilst producing Alanine (Ala). Skeletal muscle only produced Glutamine (Gln) at baseline in the volitional cycling group, otherwise the change was not significantly different from zero (Fig. 4). Neither type of exercise led to a significant change of amino acid metabolism, but it is apparent from Fig.4 that with volitional cycling there was a trend to an increase in Ala production and a decrease of glutamine production, whilst after FES cycling no such a trend was apparent (across-leg amino acid exchange remained unaffected). Uptake of BCAAs continued and did not change with either type of exercise (p=0.83 and p=0.86).

**Fig 4.**
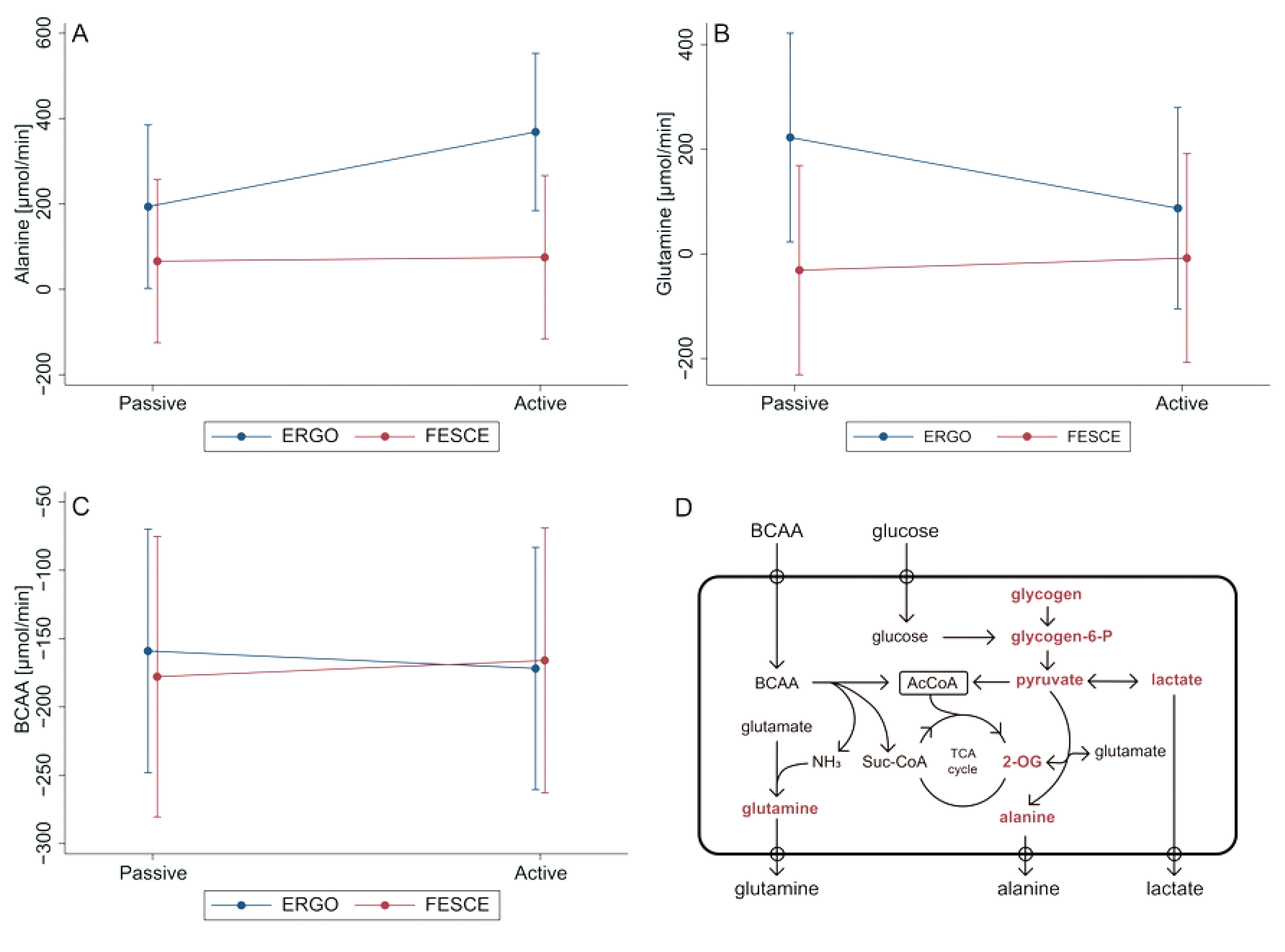
Amino acid metabolism during volitional and FES cycling. Values are derived from multiplying femoral VA differences of concentrations and calculated leg blood flow. Note: BCAA = branched-chain amino acids (i.e. the sum of Valine, Leucine, and Isoleucine); ERGO = volitional cycling; FESCE = functional electrical stimulation-assisted cycling; Passive period vs Active FES/50W volitional period. TCA = tricarboxylic acid cycle, 2-OG = 2-oxoglutarate.

## Discussion

The major finding of our study is that unloaded supine FES cycling leads to lactate production without signs of muscle hypoperfusion, as low blood flow through exercising limbs would have caused femoral venous haemoglobin desaturation (Esaki et al., 2005; Sun et al., 2016) and widening of VA-CO_2_ gap [40], which were not observed in our subjects. Moreover, there was a significant positive correlation between across-leg lactate production and femoral venous oxygenation, suggesting that subjects producing lactate did so whilst extracting less oxygen from (and producing less CO_2_ into) the local circulation. There was a marked interindividual variability in metabolic response to FES cycling: some subjects responded to FES similarly to volitional cycling, whilst others produced so much lactate that it elevated systemic (arterial) lactate concentrations well above the normal range. We have not found any convincing characteristics of the subjects producing lactate during FES, although they seemed to be leaner and oxidizing more carbohydrates at baseline. The smaller distance between skin electrodes and the muscle in lactate producers might played a role, but notably there was no correlation between the amplitude of stimulation current used and the production of lactate.

Tissue dysoxia and femoral venous desaturations are known to accompany lactate production during high intensity volitional exercise (i.e. > approx. 60% VO_2 MAX_) (Esaki et al., 2005; Gladden, 2004; Sun et al., 2016), at which oxidative phosphorylation becomes oxygen dependent. At lower exercise intensities, there is a concomitant lactate production in fast twitch glycolytic muscle fibres and consumption in slow twitch fibres [41] and - as seen in our subjects - during a steady low intensity volitional exercise, skeletal muscle may become a net lactate consumer [42].

The most obvious explanation of FES-driven lactate production would be tissue dysoxia, occurring despite adequate flow of oxygenated blood through major vessels. Non-physiological asynchronous contractions of large muscle units activated by FES [2,23] could have caused an inhomogeneous perfusion at the level of microcirculation, with hypoxic regions and units with luxurious perfusion acting as functional AV shunts. The increase in whole-body RQ with FES cycling, would support the presence of some degree of anaerobic metabolism, but it could also be explained by impaired fatty acid oxidation with the preference of carbohydrate substrates [37] or by primary increased ventilation. The major argument against microcirculatory impairment and anaerobic lactate generation is the absence of widening of venous-arterial CO_2_ gap. Carbon dioxide is produced also anaerobically and released from bicarbonate as the consequence of buffering acid load in hypoxic tissue, and because CO_2_ diffuses rapidly even from poorly perfused tissue, VA-CO_2_ gap is regarded as a very sensitive marker of tissue hypoxia caused by impaired microvascular flow [43]. Not only VA CO_2_ gap was not widened after FES cycling, but in was inversely proportional to lactate production. Moreover, the 138±29% increase in the whole body oxygen consumption after FES-cycling observed by us and others [44] would also argue against major oxygen delivery problem.

Lactate production without tissue dysoxia may occur as a result of the dysbalance between pyruvate production from glycolysis and its conversion to acetyl-CoA and oxidation in tricarboxylic acid cycle [41,42]. Muscle contraction instantly triggers, via the increase in Ca^2^+_[IC]_, glycogenolysis and glycolysis, producing pyruvate. Sudden increase in cytosolic pyruvate concentration shifts the near-equilibrium reaction: *Pyruvate* + *Glutamate* ↔ *Alanine* + *2-oxoglutarate*, rightwards. Alanine is increasingly released during exercise and 2-oxoglutarate is believed to increase the functional capacity of tricarboxylic acid cycle [45] allowing for increase in oxidative ATP production. BCAAs uptake in skeletal muscle continues or even increases during exercise, providing carbons for oxidative pathways and nitrogen for Alanine and Glutamine formation (Fig. 4D). Although non-significant, we have observed some trends to these responses after volitional cycling, but no rearrangement at all of amino acid metabolism was seen with FES exercise. Glycolytic compartment is known to respond much faster compared to oxidative phosphorylation and a rapid increase in cytosolic pyruvate concentration could lead to lactate release from cells even in the absence of tissue hypoxia [41]. Moreover, FES cycling compared to volitional exercise is known to activate glycogenolysis and glycolysis disproportionally faster than oxidative pathways [20,37]. In light of this, our data are consistent with aerobic lactate generation due to a dysbalance between pyruvate generation from glycogenolysis and glycolysis and its oxidation in citric acid cycle. Indeed, skeletal muscle is not a metabolically homogenous tissue [42] and FES may preferentially trigger muscle contraction in glycolytic fast twitch fibres, whilst lactate oxidizing slow fibres may have been less sensitive to electrical stimulation. The sensitivity of different muscle fibres to external stimulation is unknown and remains to be studied, but a higher sensitivity of fast twitch fibres would be in keeping with the finding, that a long-term external electrical stimulation of a denervated muscle restores its mass and contractile power, but not fatigability [46].

From clinical point of view we found important the absence of venous haemoglobin desaturation during FES-cycling as decreased central venous saturation impairs systemic oxygenation in patients with a degree of intrapulmonary shunt. Mild lactic acidosis could be of concern in patients with impaired lactate clearance (e.g. liver failure). Unloaded FES cycling led to VO_2_ response comparable to 25W volitional exercise, which would represent a very significant exercise load for critically ill patients, who tend to have even higher metabolic cost for a given power output [47] and only tolerated cycling at 3-6 W in one study [47]. Lastly, although the absence of laboratory signs of muscle damage and amino acid release is reassuring, the positive association of post-exercise serum myoglobin with stimulation current amplitude suggest a risk of muscle damage from the use of stimulation currents above 70mA, which are often needed to elicit visible contractions in sedated critically ill patient, perhaps due to their impaired muscle excitability [16].

The major weakness of our study is that we have not used direct measurements of leg blood flow and tissue oxygenation. However, effects of FES exercise on leg blood flow are known (Kim et al., 1995; Levine et al., 2008) and the main finding of the study, i.e. lactate production without evidence of tissue hypoxia, can be supported by across-leg VA differences alone. Muscle tissue oxygen concentrations are known to be closely reflected by femoral venous oxygen content (Mathewson et al., 2015; Sun et al., 2016).

In conclusion, we have demonstrated that 10 min of supine FES cycling in healthy volunteers leads to production of lactate without features suggestive oxygen consumption/delivery mismatch, which are known to accompany lactate production during high intensity voluntary exercise (Esaki et al., 2005; Sun et al., 2016). Despite a significant increase in systemic oxygen consumption (proportional to 25W of volitional exercise) and unaltered across-leg glucose uptake with FES cycling, we have not observed the rearrangement of amino acid metabolism towards anaplerosis.

## Acknowledgements

The authors thank to Jana Potočková, Šárka Gregorová, Šárka Vosalová for technical assistance and to all healthy volunteers, who decided to participate in this study.

## References

1. Glaser RM. Physiologic aspects of spinal cord injury and functional neuromuscular stimulation. Cent Nerv Syst Trauma. 1986;3: 49–62. Available: http://www.ncbi.nlm.nih.gov/pubmed/3524868

2. Hunt KJ,Fang J,Saengsuwan J,Grob M,Laubacher M. On the efficiency of FES cycling: a framework and systematic review. Technol Health Care. 2012;20: 395–422. doi:10.3233/THC-2012-0689

3. Szecsi J,Schiller M. FES-propelled cycling of SCI subjects with highly spastic leg musculature. NeuroRehabilitation. 2009;24: 243–53. doi:10.3233/NRE-2009-0475

4. Lo H-C, Hsu Y-C, Hsueh Y-H, Yeh C-Y. Cycling exercise with functional electrical stimulation improves postural control in stroke patients. Gait Posture. 2012;35: 506–10. doi:10.1016/j.gaitpost.2011.11.017

5. Peri E,Ambrosini E,Pedrocchi A,Ferrigno G,Nava C,Longoni V, et al. Can FES-Augmented Active Cycling Training Improve Locomotion in Post-Acute Elderly Stroke Patients? Eur J Transl Myol. PAGEPress; 2016;26: 6063. doi:10.4081/ejtm.2016.6063

6. Szecsi J,Schlick C,Schiller M,Pöllmann W,Koenig N,Straube A. Functional electrical stimulation-assisted cycling of patients with multiple sclerosis: Biomechanical and functional outcome - A pilot study. J Rehabil Med. 2009;41: 674–680. doi:10.2340/16501977-0397

7. Mohr T,Dela F,Handberg A,Biering-Sørensen F,Galbo H,Kjaer M. Insulin action and long-term electrically induced training in individuals with spinal cord injuries. Med Sci Sports Exerc. 2001;33: 1247–52. Available: http://www.ncbi.nlm.nih.gov/pubmed/11474322

8. Young W. Electrical Stimulation and Motor Recovery. Cell Transplant. 2015;24: 429–446. doi:10.3727/096368915X686904

9. Morris PE,Herridge MS. Early intensive care unit mobility: future directions. Crit Care Clin. Elsevier; 2007;23: 97–110. doi:10.1016/j.ccc.2006.11.010

10. Schweickert WD,Kress JP. Implementing Early Mobilization Interventions in Mechanically Ventilated Patients in the ICU. Chest. 2011;140: 1612–1617. doi:10.1378/chest.10-2829

11. Choong K,Koo KKY,Clark H,Chu R,Thabane L,Burns KEA,et al. Early Mobilization in Critically Ill Children. Crit Care Med. 2013;41: 1745–1753. doi:10.1097/CCM.0b013e318287f592

12. TEAM Study Investigators,Hodgson C,Bellomo R,Berney S,Bailey M,Buhr H,et al. Early mobilization and recovery in mechanically ventilated patients in the ICU: a bi-national, multi-centre, prospective cohort study. Crit Care. 2015;19: 81. doi:10.1186/s13054-015-0765-4

13. Pawlik AJ. Early Mobilization in the Management of Critical Illness. Crit Care Nurs Clin North Am. 2012;24: 481–490. doi:10.1016/j.ccell.2012.05.003

14. Friedrich O,Reid MB,Van den Berghe G,Vanhorebeek I,Hermans G,Rich MM,et al. The Sick and the Weak: Neuropathies/Myopathies in the Critically Ill. Physiol Rev. American Physiological Society; 2015;95: 1025–109. doi:10.1152/physrev.00028.2014

15. Herridge MS,Tansey CM,Matté A,Tomlinson G,Diaz-Granados N,Cooper A,et al. Functional Disability 5 Years after Acute Respiratory Distress Syndrome. N Engl J Med. 2011;364: 1293–1304. doi:10.1056/NEJMoa1011802

16. Kress JP,Hall JB. ICU-Acquired Weakness and Recovery from Critical Illness. N Engl J Med. 2014;370: 1626–1635. doi:10.1056/NEJMra1209390

17. Levine S,Nguyen T,Taylor N,Friscia ME,Budak MT,Rothenberg P,et al. Rapid Disuse Atrophy of Diaphragm Fibers in Mechanically Ventilated Humans. N Engl J Med. 2008;358: 1327–1335. doi:10.1056/NEJMoa070447

18. Parry SM,Puthucheary ZA. The impact of extended bed rest on the musculoskeletal system in the critical care environment. Extrem Physiol Med. BioMed Central; 2015;4: 16. doi:10.1186/s13728-015-0036-7

19. Parry SM,Berney S,Warrillow S,El-Ansary D,Bryant AL,Hart N,et al. Functional electrical stimulation with cycling in the critically ill: A pilot case-matched control study. J Crit Care. 2014;29: 695.e1–695.e7. doi:10.1016/j.jcrc.2014.03.017

20. Duffell LD,de N Donaldson N,Newham DJ. Why is the Metabolic Efficiency of FES Cycling Low? IEEE Trans Neural Syst Rehabil Eng. 2009;17: 263–269. doi:10.1109/TNSRE.2009.2016199

21. Hunt KJ,Hosmann D,Grob M,Saengsuwan J. Metabolic efficiency of volitional and electrically stimulated cycling in able-bodied subjects. Med Eng Phys. 2013;35: 919–925. doi:10.1016/j.medengphy.2012.08.023

22. Hunt KJ,Ferrario C,Grant S,Stone B,McLean AN,Fraser MH,et al. Comparison of stimulation patterns for FES-cycling using measures of oxygen cost and stimulation cost. Med Eng Phys. Elsevier; 2006;28: 710–8. doi:10.1016/j.medengphy.2005.10.006

23. Downey RJ,Merad M,Gonzalez EJ,Dixon WE. The Time-Varying Nature of Electromechanical Delay and Muscle Control Effectiveness in Response to Stimulation-Induced Fatigue. IEEE Trans Neural Syst Rehabil Eng. 2017;25: 1397–1408. doi:10.1109/TNSRE.2016.2626471

24. Kjaer M,Perko G,Secher NH,Boushel R,Beyer N,Pollack S,et al. Cardiovascular and ventilatory responses to electrically induced cycling with complete epidural anaesthesia in humans. Acta Physiol Scand. Blackwell Publishing Ltd; 1994;151: 199–207. doi:10.1111/j.1748-1716.1994.tb09738.x

25. Kim CK,Strange S,Bangsbo J,Saltin B. Skeletal muscle perfusion in electrically induced dynamic exercise in humans. Acta Physiol Scand. Blackwell Publishing Ltd; 1995;153: 279–287. doi:10.1111/j.1748-1716.1995.tb09864.x

26. Scremin OU,Cuevas-Trisan RL,Scremin AM,Brown C V,Mandelkern MA. Functional electrical stimulation effect on skeletal muscle blood flow measured with H2(15)O positron emission tomography. Arch Phys Med Rehabil. 1998;79: 641–6. Available: http://www.ncbi.nlm.nih.gov/pubmed/9630142

27. Tepavac D,Schwirtlich L. Detection and prediction of FES-induced fatigue. J Electromyogr Kinesiol. 1997;7: 39–50. Available: http://www.ncbi.nlm.nih.gov/pubmed/20719690

28. Kim CK,Bangsbo J,Strange S,Karpakka J,Saltin B. Metabolic response and muscle glycogen depletion pattern during prolonged electrically induced dynamic exercise in man. Scand J Rehabil Med. 1995;27: 51–8. Available: http://www.ncbi.nlm.nih.gov/pubmed/7792551

29. Glaser RM. Physiology of Functional Electrical Stimulation-Induced Exercise: Basic Science Perspective. Neurorehabil Neural Repair. Sage PublicationsSage CA: Thousand Oaks, CA; 1991;5: 49–61. doi:10.1177/136140969100500106

30. Dreyer HC,Fujita S,Cadenas JG,Chinkes DL,Volpi E,Rasmussen BB. Resistance exercise increases AMPK activity and reduces 4E-BP1 phosphorylation and protein synthesis in human skeletal muscle. J Physiol. Wiley-Blackwell; 2006;576: 613–24. doi:10.1113/jphysiol.2006.113175

31. Hulston CJ,Wolsk E,Grondhal TS,Yfanti C,Van Hall G. Protein Intake Does Not Increase Vastus Lateralis Muscle Protein Synthesis during Cycling. Med Sci Sport Exerc. 2011;43: 1635–1642. doi:10.1249/MSS.0b013e31821661ab

32. Biolo G,Maggi SP,Williams BD,Tipton KD,Wolfe RR. Increased rates of muscle protein turnover and amino acid transport after resistance exercise in humans. Am J Physiol Metab. 1995;268: E514–E520. doi:10.1152/ajpendo.1995.268.3.E514

33. Holm L,van Hall G,Rose AJ,Miller BF,Doessing S,Richter EA,et al. Contraction intensity and feeding affect collagen and myofibrillar protein synthesis rates differently in human skeletal muscle. Am J Physiol Metab. 2010;298: E257–E269. doi:10.1152/ajpendo.00609.2009

34. Dideriksen K,Reitelseder S,Holm L. Influence of Amino Acids, Dietary Protein,and Physical Activity on Muscle Mass Development in Humans. Nutrients. 2013;5: 852–876. doi:10.3390/nu5030852

35. van Hall G,González-Alonso J,Sacchetti M,Saltin B. Skeletal muscle substrate metabolism during exercise: methodological considerations. Proc Nutr Soc. 1999;58: 899–912. Available: http://www.ncbi.nlm.nih.gov/pubmed/10817157

36. Tuma P. Rapid determination of globin chains in red blood cells by capillary electrophoresis using INSTCoated fused-silica capillary. J Sep Sci. 2014;37: 1026–1032. Available: http://dx.doi.org/10.1002/jssc.201400044

37. Kjær M,Dela F,Sørensen FB,Secher NH,Bangsbo J,Mohr T,et al. Fatty acid kinetics and carbohydrate metabolism during electrical exercise in spinal cord-injured humans. Am J Physiol Integr Comp Physiol. American Physiological SocietyBethesda,MD; 2001;281: R1492–R1498. doi:10.1152/ajpregu.2001.281.5.R1492

38. Esaki K,Hamaoka T,Rådegran G,Boushel R,Hansen J,Katsumura T,et al. Association between regional quadriceps oxygenation and blood oxygen saturation during normoxic one-legged dynamic knee extension. Eur J Appl Physiol. 2005;95: 361–370. doi:10.1007/s00421-005-0008-5

39. Sun Y,Ferguson BS,Rogatzki MJ,McDonald JR,Gladden LB. Muscle Near-Infrared Spectroscopy Signals versus Venous Blood Hemoglobin Oxygen Saturation in Skeletal Muscle. Med Sci Sport Exerc. 2016;48: 2013–2020. doi:10.1249/MSS.0000000000001001

40. Vallet B,Teboul J-L, Cain S,Curtis S. Venoarterial CO(2) difference during regional ischemic or hypoxic hypoxia. J Appl Physiol. 2000;89: 1317–1321. doi:10.1152/jappl.2000.89.4.1317

41. Gladden LB. Lactate metabolism: a new paradigm for the third millennium. J Physiol. Wiley-Blackwell; 2004;558: 5–30. doi:10.1113/jphysiol.2003.058701

42. Brooks GA. Intra- and extra-cellular lactate shuttles. Med Sci Sports Exerc. 2000;32: 790–9. Available: http://www.ncbi.nlm.nih.gov/pubmed/10776898

43. Mallat J,Lemyze M,Meddour M,Pepy F,Gasan G,Barrailler S,et al. Ratios of central venous-to-arterial carbon dioxide content or tension to arteriovenous oxygen content are better markers of global anaerobic metabolism than lactate in septic shock patients. Ann Intensive Care. Springer; 2016;6: 10. doi:10.1186/s13613-016-0110-3

44. Hettinga DM,Andrews BJ. Oxygen consumption during functional electrical stimulation-assisted exercise in persons with spinal cord injury: implications for fitness and health. Sports Med. 2008;38: 825–38. Available: http://www.ncbi.nlm.nih.gov/pubmed/18803435

45. Wagenmakers AJ. Muscle amino acid metabolism at rest and during exercise: role in human physiology and metabolism. Exerc Sport Sci Rev. 1998;26: 287–314. Available: http://www.ncbi.nlm.nih.gov/pubmed/9696993

46. Ashley Z,Sutherland H,Russold MF,Lanmüller H,Mayr W,Jarvis JC,et al. Therapeutic stimulation of denervated muscles: The influence of pattern. Muscle Nerve. 2008;38: 875–886. doi:10.1002/mus.21020

47. Hickmann CE,Roeseler J,Castanares-Zapatero D,Herrera EI,Mongodin A,Laterre P-F. Energy expenditure in the critically ill performing early physical therapy. Intensive Care Med. 2014;40: 548–555. doi:10.1007/s00134-014-3218-7

48. Mathewson KW,Haykowsky MJ,Thompson RB. Feasibility and reproducibility of measurement of whole muscle blood flow, oxygen extraction,and VO 2 with dynamic exercise using MRI. Magn Reson Med. 2015;74: 1640–1651. doi:10.1002/mrm.25564

